# Turmeric shortens lifespan in houseflies

**DOI:** 10.1101/2024.01.07.574512

**Authors:** Sophie Laurie, Leah Ainslie, Sharon Mitchell, Juliano Morimoto

## Abstract

Climate change poses a significant threat to food security and global public health with the increasing likelihood of insect pest outbreaks. Alternative ways to control insect populations, preferably using environmental-friendly compounds, are needed. Turmeric has been suggested as a natural insecticide with toxicity properties in some insect groups. However, empirical evidence of the effects turmeric – and their interaction with other ecological factors such as diet – on insect survival has been limited. Here, we tested the effects of turmeric and its interactions with diets differing in protein source in the common housefly, *Musca domestica*. We found that turmeric shortened lifespan independent of diet and sex. Females in turmeric diets were heavier at death, which was likely driven by a combination of relatively lower rates of body mass loss during their lifetime and a higher percentage of water content at death. Each sex responded differently to the protein source in the diet, and the magnitude of the difference in lifespan between sexes were greatest in diets in which protein source was hydrolysed yeast; individuals from both sexes lived longest in sucrose-milk diets and shortest in diets with hydrolysed yeast. There was no evidence of an interaction between turmeric and diet, suggesting that the toxicity effects are independent of protein source in the diet. Given the seemingly opposing effects of turmeric in insects and mammals being uncovered in the literature, our findings provide further evidence in support of turmeric as a potential natural insecticide.

## Introduction

Insect pests inflict damage to agricultural crops resulting in yield loss which causes major economic losses worldwide. This damage is predicted to increase with climate change (Gomez-Zavaglia et al., 2020; Tonnang et al., 2022), as insects are sensitive to the rise in global temperatures and will pose further threats to food security (Lehmann et al., 2020; Skendžić et al., 2021). Pesticides are chemicals used to control insect outbreaks and have been extensively used in agriculture. Approximately two million tonnes of pesticides are used worldwide per year, which pollute ecosystems and pose serious health hazards for living organisms (Sharma et al., 2019; Damalas & Eleftherohorinos, 2011). For instance, dichlorodiphenyltrichloroethane (DDT) can remain in soils and water for years after exposure posing health risks for humans and other living organisms (Beard, 2006; Turusov et al., 2002). Alternative ways to control insect pest outbreaks are needed, which are affordable and more environmental-friendly.

Natural compounds can have pesticide activity, making them attractive for further studies as environmental-friendly alternatives to chemical pesticides (Batish et al., 2008: Duke et al., 1990; Schmutterer, 1990). Turmeric (*Curcuma longa*) is a spice that is native to southeast Asia and is used worldwide as a medicinal plant, food colouring agent, and an ingredient in supplements (Gupta et al., 2013; Iweala et al., 2023). Studies have shown that turmeric is toxic to insects, making this compound a candidate as natural pest control agent (Hellfeld et al., 2023; Raje et al., 2015; Uysal et al., 2015; Damalas, 2011, De Souza Tavares et al., 2016). For example, a study by Muhammad et al., (2018) found turmeric extracts to be effective against insect pests that were feeding on okra (*Abelmoschus esculentus*). The results showed turmeric significantly protected the okra plant from aphid (*Aphis gossypii*) attacks and was more effective than synthetic pesticides that had been previously used. Turmeric extracts have also shown to reduce the development of the red flour beetle *Tribollium castaneum*, which helps to stop the invasion of the beetle against wheat grains (Ali et al., 2014). Similarly, a study by De Souza Tavares et al., (2016) extracted *ar*-turmerone from *C. longa* rhizomes to investigate its insecticidal effects against the cabbage looper. This study found *ar*-turmerone reduced larval weight and increased mortality of the cabbage looper which helped to stop the insect from feeding on agricultural crops. Other studies have used turmeric oils as a natural and effective insecticide to repel the grain borer pest *Rhyzopertha dominica* from feeding on agricultural crops (Visakh et al., 2023; Jilani & Saxena, 1990). These oils have been described as having insect repellent properties and have been used as an insecticide against other indoor pests including the acid flies *Paederus fuscipes* and the darkling beetles *Luprops tristis* (Vineesh et al., 2023). These essential oils from turmeric also show promising repellent activity against mosquito species (Damalas, 2011). Turmeric crude oil extract can also help to control cucumber powdery mildew which is a toxic foliar disease caused by the plant pathogen *Podosphaera xanthii.* This disease damages the yield and reduces the quality of crops, but turmeric oils show successful fungicidal effects against *P*. *xanthii* (Fu et al., 2018). Turmeric is relatively cheap and more environmentally friendly alternative to chemical pesticides. Thus, turmeric use as pesticide can contribute towards more sustainable crop production that minimizes environmental contamination and human health risks (Roy et al., 2014).

Nutrition is essential for insect growth and development but can also act as – or modulate the effects of – toxic compounds (Deans et al., 2017; Wada-Katsumata et al., 2021). For example, recent studies have shown that excess protein can be toxic and shorten adult lifespan in flies and crickets (Lee et al., 2008; Jensen et al., 2015; Fanson et al., 2009; Fanson et al., 2012; Prabhu et al., 2008; Maklakov et al., 2008). Other studies have also shown that protein shortens lifespan in different insect species including ants (Dussutour & Simpson, 2012; Dussutour et al., 2016; Kay et al., 2010), caterpillars (Despland & Noseworthy, 2006) and honeybees (Pirk et al., 2010). Diet also modulates insect survival and behaviour by playing an interactive role with toxic bait compounds. For example, in the German cockroach *Blattella germanica,* both protein-to-carbohydrate ratios (PC ratio) and sugar type (glucose versus fructose) interacted with the formulation of the bait containing insecticide hydramethylnon to reduce survival of adult males and first-instar nymphs (Ko et al., 2016). In fact, exposure to suboptimal, high-protein diets were improved efficacy of bait combinations, suggesting that a high-protein diet could potentialize the toxicity of the bait. Notably though, this study investigates the interaction of diet and a noxious compound in the context of baits. Recent studies have broadened this approach and investigated the interactive effects of diet and toxic natural compounds. For example, Morimoto (2022) showed that urea was toxic for the development of *Drosophila melanogaster* larvae but had no effect on oviposition choices by egg-laying females. Moreover, it showed that high-sugar diets combined with low and intermediate concentrations of urea were potentially more detrimental to the development of the larvae than high-protein diets with similar urea concentrations (Morimoto, 2022). Collectively, these results suggest that (1) diet is a major ecological factor determining development and health of insects, (2) diet modulates the toxicity effects of chemical baits as well as (3) natural toxic compounds. To date, however, we have very little understanding of how diet and natural toxic compounds interact, most of which comes from the work in model species such as *D. melanogaster* (see e.g., Hellfeld et al., 2023; Raje et al., 2015; Uysal et al., 2015; Damalas, 2011, De Souza Tavares et al., 2016).

To address this, we manipulated protein source in the diet and turmeric levels to investigate how the toxicity of turmeric interacts with diet to modulate housefly *Musca domestica* lifespan and body composition. The common housefly, *Musca domestica,* is a cosmopolitan species closely associated with humans, and is perhaps one of the most common insect disease vectors of our societies (Cervelin et al., 2018; Neupane et al., 2019; Poudel et al., 2019; Raele et al., 2021; Toto et al., 2022). With our changing climate, houseflies could become an increasing threat to public health through the increase risks of disease transmission (Cousins et al., 2019; Meraz Jiménez et al., 2019). Here, we tested whether turmeric interacted with diet to shorten male and female lifespan, and then investigated individuals’ body mass at death, loss of body mass over lifespan and body water content at death. We predicted that turmeric would shorten individual lifespan due its known toxic effects. We also predicted that flies feeding on sucrose alone would have a longer lifespan, based on the findings in the literature for other insect species (Fanson et al., 2009; Lee et al., 2008). There was not enough *a priori* information to conceptualise predictions about the interaction between turmeric and diet. We predicted that body mass at death, weight loss, and water content would be lower in flies in turmeric diets, likely because of feeding avoidance due to the presence of turmeric. Our results reveal how a noxious compound and diet can be used to modulate lifespan and body traits in a cosmopolitan insect species of public health interest.

## Material and methods

### Fly stock and experimental flies

Houseflies were maintained in a large stock population (>700 individuals) at 25°C and 35% humidity, with a 12:12 hour light:dark cycle. Flies were obtained as pupae from a commercial supplier (Blades Biologics Ltd) and were maintained in the laboratory for 5 generations prior to the experiments. Stock colonies were maintained inside mesh cages (32cmx32cmx32cm) (BugDorm-4E4545; MegaView science Co. Limited). Adults were given *ad libitum* access to Hydrolysed Yeast (MP Biomedicals, Cat no: 103304) and commercial sucrose (Tate & Lyle White Granulated Sugar). Water was provided using a 500ml lidded Plastic Container with a slit cut into the centre of the lid which gave access to water through a moist surface and minimised drowning. We collected eggs to generate experimental flies by introducing a plastic container (14cm x 9cm x 5cm) into the stock cage containing the larval diet used to maintain our stocks. Larval diet recipe included 70g Organic Wheat Bran, 6g Nestle Instant Full Cream Milk Powder, 15g Alfalfa meal from Supreme Science Selective Guinea Pig Pellets (blended) and 300ml water. Females were allowed to lay eggs freely for 12 hours, after which containers were placed onto 1-2 cm of ground wheat bran within a larger 3 litre container (23cm x 16cm x 8cm) and reserved at controlled temperature (see above) to complete development. Larval diet was monitored daily, and water was added to the container to avoid desiccation. After 10 days, the larval diet was flooded with approximately 200ml of water to encourage pupation. Pupae for the experiments were retrieved from bran using a commercial sieve, reserved in Petri dishes at controlled temperatures for 7 days, and assigned to one of the experimental diets.

### Experimental diets and turmeric

Pupae were weighted (see details below) and randomly allocated to plastic vials (SARSTEDT, Ref: 58.631, 75 x 23.5mm) containing 5mL of each of the four experimental diets (henceforth ‘diets’) which varied in the protein source that was available to the flies: Sucrose, Brewer’s Yeast (BY) (MP Biomedicals, Cat no: 903312), hydrolysed yeast (HY) (MP Biomedicals, Cat no: 103304) and whole milk (Nestle Instant Full Cream Milk Powder, Nestlé) (M) diets. From manufacturer specifications, HY was the diet with the highest protein content (ca. 60%), followed by BY (ca 40%) and M (ca. 25.7%). Table 1 summarises the diet recipes. Each diet had two turmeric levels: control (0%) and treatment (5%) turmeric (w/v) (Turmeric from Sevenhills Wholefoods®).

### Survival and body composition

We investigated how adult diet composition and the presence of turmeric affected survival time and body composition. We randomly sampled 280 pupae collected as described above and placed each pupa in one of the experimental treatments (N_total_ = 280). Adults in the experimental treatments were maintained in isolation for the entire experiment and were monitored daily which allowed us to measure lifespan. Once the individuals died, they were frozen at -20°C within 24 h. We then processed the individuals by measuring their weight at death and water content at death. Water content at death was obtained by subtracting the weight at death from the weight after individuals were maintained at 65°C for 2 days; water content at death was the different between wet body weight at death minus dry body weight at death, divided by wet body weight at death x 100 (for percentage). All individuals were weighed in a SECURA124-1S, Secura® Laboratory Balances, Sartorius (precision: 0.00001g).

### Statistical analysis

All statistical analyses were conducted in RStudio version 4.2.2. We used the ‘ggplot2’ package version 3.4.3 (Wickham, 2016) for data visualisation (Wickham et al., 2019). We wrangled the data using ‘stringr’ version 1.5.0 (Wickham, 2022); ‘car’ version 4.0.0 (Fox & Weisberg, 2019); ‘dplyr’ version 1.1.3 (Wickham, François, et al., 2023); ‘survival’ version 3.5-7 (Therneau, 2023); ‘ggfortify’ version 0.4.16 (Horikoshi & Tang, 2018) and ‘tidyr’ packages version 2.0.0 (Wickham, Vaughan, et al., 2023). We fitted a survival model with Weibull distribution to assess the effects of diet, turmeric, and sex on fly survival. We fitted linear models with either adult weight at death, weight loss from pupae to adult at death, and water content at death as response variables, and the main, two-way and three-way interactions of turmeric, diet and sex as explanatory variables. To improve model fit, we transformed adult weight at death (weight at death^2) and water content (square-root). P-values were obtained from F-statistics using the inbuilt anova function in R for all models except survival, the latter which p-values were obtained from a X^2^log-rank test using the ‘Anova’ function of the ‘car’ package.

## Results

### Turmeric shortens lifespan in a diet- and sex-independent fashion

We first tested whether turmeric addition to diets with varying protein sources shortened individual lifespan. There was a statistically significant two-way interaction between diet and sex (X^2^ = 62.849, p < 0.001). This was driven by the fact that males had substantially shorter lifespan than females when the protein source was hydrolysed yeast (HY), but this effect disappeared in sucrose only diets and diets with brewers’ yeast (BY) or whole milk (M) as protein sources (Fig 1a). There were also statistically significant effects of diet (X^2^ = 105.318, p < 0.001), turmeric (X^2^ = 15.884, p < 0.001) and sex (X^2^ = 16.949, p < 0.001). These effects were driven by the fact that, on average, turmeric addition to the diet or HY as protein source shortened lifespan of both males and females, whereas males were on average shorter lived than females (Fig 1a; Table S1). The three-way interaction between diet, turmeric, and sex was not statistically significant (X^2^ = 6.347, p = 0.096), neither were the two-way interactions between turmeric and sex (X^2^ = 0.026, p = 0.872) and diet and turmeric (X^2^ = 6.412, p = 0.093). These results showed that turmeric addition shortened lifespan in a diet- and sex-independent fashion, corroborating the potential role of turmeric as noxious compound to shorten insect lifespan. Moreover, the results also showed that protein source in the diet can have sex-specific effects on lifespan above and beyond the presence of turmeric.

**Figure 1.**
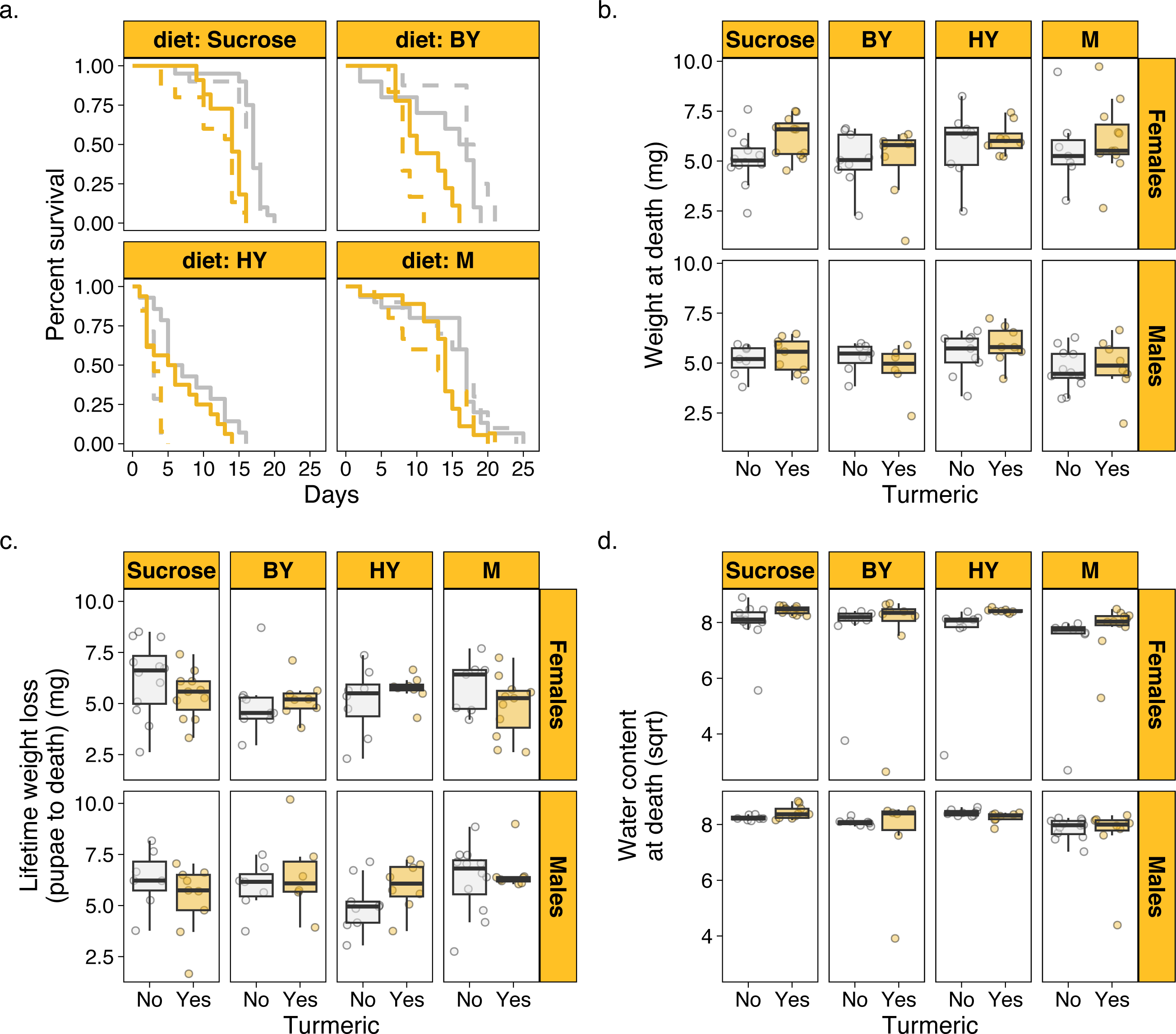
Turmeric shortens lifespan and affect body traits in houseflies. (a) Female (solid) and male (dashed) survival across diets varying in protein source with (yellow) and without turmeric (grey). (b) Body weight at death across diets with and without turmeric. (c) Lifetime weight loss (from pupae to adult death) across diets with and without turmeric. (d) Water content at death (square root-transformed) across diets with and without turmeric. M: Whole Milk as protein source; BY: Brewers’ yeast as protein source; HY: Hydrolysed yeast as protein source.

### Turmeric and diet modulate body traits, although not necessarily interactively

Next, we investigated the effects of turmeric and protein source in the diets on organismal traits such as body weight at death and water content at death, which were used here as proxies for individual condition. After controlling for differential survival across treatments, there was a statistically significant effect of sex (F_1,123_ = 7.159, p = 0.008) and turmeric (F_1,123_ = 6.536, p = 0.011), whereby female flies on turmeric diets were on average heavier at death (Fig 1b). This effect could have emerged from two factors. First, female flies on turmeric diet could be heavier because they lost relatively less weight over their lifespan. Indeed, after controlling for differential survival, males lost more weight from pupae to adult death (F_1,123_ = 5.163, p = 0.024), but turmeric had no effect on weight loss (F_1,123_ = 0.035, p = 0.851; Table S1; Fig 1c). As a result, weight loss alone could not fully explain the results the effects of turmeric and sex on adult weight at death. A second explanation for the effects observed for weight at death could be due to differential water loss. Insect adult body composition is primarily related to water content (Molleman et al., 2001; Fairbanks & Burch, 1970). Thus, turmeric could have affected individual hydration, which in turn influenced their weight at death. After controlling for differential survival, there was a statistically significant interaction between turmeric and sex (F_1,123_ = 3.983, p = 0.048; Table S1), which was driven primarily by the fact that females on turmeric diets had on average higher water content at death (Fig 1d). There was also a small but statistically significant effect of diet (F_1,123_ = 3.055, p = 0.0310), driven by lower water content at death in M diets for both sexes and irrespective of turmeric (Fig 1d). These results showed that body weight at death was higher for females in turmeric diets likely due to a combination of lower body weight loss over lifespan and a relatively higher water content at death for females in turmeric diets. There were no other statistically significant three-way, two-way or main effects in the models of weight loss and water content (Table S1).

## Discussion

We set out to test the additive and interactive effects of turmeric in diets with varying protein sources in houseflies. Our data showed that turmeric shortened lifespan in a diet- and sex-independent fashion. This confirms our prediction that turmeric is a toxic compound which could potentially be used to control the development of houseflies. We also found that turmeric and diet independently modulated organism traits such as body mass loss during their lifetime and a higher percentage of water content at death. There was no evidence that turmeric interacted with diet. We also found that females had longer lifespan than males across all diets with and without turmeric, but the magnitude of the difference was diet-dependent but turmeric-independent. Together, our findings show that a natural compound such as turmeric can be used as an insecticide in a cosmopolitan fly species.

Our findings corroborate the role of turmeric in shortening lifespan from previous studies (Fig 1a). For instance, Uysal et al., (2015) found that turmeric shortened lifespan in *D. melanogaster* adults. Similarly, Hellfeld et al., (2023) found that increasing turmeric concentration in the diet decreased oviposition in adult females and subsequently reduced egg-to-adult viability in the developing larvae in *D. malanogaster.* In fact, the study showed that high turmeric levels (>5%) could lead to complete developmental arrest (Hellfeld et al., 2023). The molecular mechanisms through which turmeric – or compounds therein – leads to toxicity in insects is uncertain but Rahman et al., (2022) found that higher concentrations of turmeric (>1%) in the diet decreased β-tubulin levels in the brain and affected a range of physiological traits of *D. melanogaster.* The study also found that an optimal dosage of 0.5% of turmeric could maintain healthy aging by increasing β-tubulin expression. Moreover, 0.5% turmeric also improved survivability, locomotor activity, fertility, tolerance to oxidative stress, and eye health (Rahman et al., 2022). Taken together, these results suggest that turmeric has a non-linear effect on insect health: at lower concentrations, turmeric might be beneficial by increasing β-tubulin and protecting the physiology of the individual up to a point, after which turmeric is toxic and leads to negative effects that decrease survival. This leads us to hypothesise that the relationship between turmeric concertation in the diet and β- tubulin expression in the brain is a concave parabola. Having said that, Hellfedt et al. (2023) conducted a discrete dose-response study and found that turmeric was harmful for the developing larvae at concentrations above 1% but found no evidence of beneficial effects. Dose-response studies for turmeric in other species are needed to uncover the nuances of the toxic vs beneficial effects of turmeric in insects.

Curcumin, one of the main components of turmeric, was found to extend the lifespan in *D. melanogaster* (Suckow & Suckow, 2006, Chen et al., 2018, Shen et al., 2013). Curcumin prolongs lifespan in *D. melanogaster* by enhancing superoxide dismutase (SOD) activity. Similarly, Chen et al., (2018) also found that under heat stress conditions, adding curcumin to diet eased the effects of increased oxidative stress by increasing SOD expression and resulting in longer lifespan. Shen et al., (2013) found that *D. melanogaster* fed diets with either 0.5 or 1.0mg/g of curcumin in the diet also increased the enzyme activity of SOD in both adult male and females. Curcumin extended the lifespan and improved the physiological traits that are related to aging. It is therefore possible that turmeric effects on survival are modulated directly by curcumin, and that curcumin might itself be beneficial while other compounds in turmeric may be triggering toxicity. The benefits of curcumin are likely observed due to its antioxidant role. It is unclear why an excess of curcumin could lead to toxicity, and future studies should conduct a dose-response experiment with both curcumin and turmeric to disentangle the roles of each of these compounds in the beneficial vs toxic effects in insects.

Our findings that that females on turmeric diets were on average heavier at death contradicts a previous study by Rawal et al., (2014) which showed that flies feeding on turmeric had no noticeable effect on body weight in *D. melanogaster*, although there was a slight decline in body weight in their experiment which was attributed to the process of aging. More studies are needed to corroborate whether turmeric feeding affect body traits in insects. Nevertheless, our data supports previous findings in nutritional research and showed that flies feeding on protein-rich diets had shortened lifespan. This is because amongst the protein sources used here, hydrolysed yeast contained the highest protein concentration (∼60% according to manufacturer report). This corroborates previous findings in other flies that protein-rich diets shortened lifespan (Lee et al., 2008; Jensen et al., 2015; Fanson et al., 2009; Fanson et al., 2012; Prabhu et al., 2008), which has also been found in other insect species [e.g. (Maklakov et al., 2008; Dussutour & Simpson, 2012; Dussutour et al., 2016; Kay et al., 2010; Despland & Noseworthy, 2006; Pirk et al., 2010)]. In fact, a recent comparative nutrition study showed that lifespan is maximized at lower protein-to-carbohydrate ratios, and that Dipterans often survive the longest in sugar-rich diets (Morimoto et al., 2023). Overall, our results add to this body of literature by showing that houseflies also survive longer in sugar-rich diets.

## Conclusion

Our study shows that turmeric can be an important ally to shorten lifespan of houseflies, a species with public health interest. This adds to the growing literature showing that turmeric can be used to manage insect pests of agriculture. Notably, turmeric has known health benefits to mammals and humans [e.g. (Labban et al., 2004; Bhat et al., 2019)]. This makes turmeric the perfect candidate to be used in agricultural insect pest control because it can have no – or even positive – effects on animals and humans that consume the food treated with turmeric. We urge future studies to focus attention on two topics, should turmeric be considered as natural insecticide. First, studies should focus on better understanding the underlying reasons as to why turmeric is toxic in insects but appear beneficial in mammals and humans. We hypothesise that insects have a lower threshold tolerance for turmeric concentrations compared with mammals and humans, which would explain why toxicity effects are observed primarily in the former but not in the latter groups. Second, studies should focus on modelling the feasibility of using turmeric in large-scale agricultural settings, and its effectiveness on controlling pest outbreak. Overall, turmeric is emerging as a potent natural pesticide in both fundamental and applied entomology and our study corroborates the potential uses of turmeric for insect control.

## Acknowledgements

The authors would like to thank Elissavet Kaplaneli for the technical support in keeping the housefly colonies.

## Authors’ contributions

SL and JM designed the study. SL collected the data. SL and JM analysed the data and wrote the manuscript. All authors contributed to the writing of the final version of the manuscript and approved its submission.

## Data availability statement

Data will be available in Dryad upon acceptance of the manuscript.

## Supplementary Material

Table S1. Complete model output for the analysis presented in the main text.

